# Interpretable and predictive models based on high-dimensional data in ecology and evolution

**DOI:** 10.1101/2024.03.15.585297

**Authors:** Joshua P. Jahner, C. Alex Buerkle, Dustin G. Gannon, Eliza M. Grames, S. Eryn McFarlane, Andrew Siefert, Katherine L. Bell, Victoria L. DeLeo, Matthew L. Forister, Joshua G. Harrison, Daniel C. Laughlin, Amy C. Patterson, Breanna F. Powers, Chhaya M. Werner, Isabella A. Oleksy

## Abstract

The proliferation of high-dimensional data in ecology and evolutionary biology raises the promise of statistical and machine learning models that are highly predictive and interpretable. However, high-dimensional data are commonly burdened with an inherent trade-off: in-sample prediction of outcomes will improve as additional variables are included in the model, but this may come at the cost of poor predictive accuracy and limited generalizability for future or unsampled observations (out-of-sample prediction). To confront this problem of overfitting, sparse models can focus on key variables by correctly placing low weight on unimportant variables. We competed nine methods to quantify their performance in variable selection and prediction using simulated data with different sample sizes, numbers of variables, and strengths of effects. Overfitting was typical for many methods and simulation scenarios. Despite this, in-sample and out-of-sample prediction converged on the true predictive target for simulations with more observations, larger causal effects, and fewer variables. Accurate variable selection to support process-based understanding will be unattainable for many realistic sampling schemes in ecology and evolution. We use our analyses to characterize data attributes for which statistical learning is possible, and illustrate how some sparse methods can achieve predictive accuracy while mitigating and learning the extent of overfitting.

## 1 Introduction

Research in ecology and evolution has seen dramatic growth in data due to technological advances for automation and high-throughput sampling (e.g., water or air sampling, Porter *et al*. 2009; satellite imagery, Ustin & Middleton 2021, Cavender-Bares *et al*. 2022; DNA sequencing, Halldors-son *et al*. 2022, Rubinacci *et al*. 2023; and GPS telemetry, Wilmers *et al*. 2015, Gigliotti *et al*. 2022). While large data sets (i.e. many records (N), many predictor variables (P), or both) have the potential to greatly improve our understanding of complex systems across the continuum of modeling purposes (e.g. data exploration, inference, prediction; Dormann *et al*. 2012, Tredennick *et al*. 2021), they also pose considerable challenges for data analysis and incorporation into formal process models. For example, a cross-cutting objective in ecology and evolutionary biology involves learning the causes of species abundances and distributions, including making predictions about the responses of wild and cultivated organisms to climate change (Faske *et al*. 2023, Forister *et al*. 2023, Grames & Forister 2024, Halsch *et al*. 2024, Laughlin & McGill 2024, Li *et al*. 2024). These predictions are commonly based on contemporary observations of organisms and many dimensions of abiotic and biotic environments, and intrinsic attributes (e.g., the genome) that could be causally associated with their species distributions (e.g., Faske *et al*. 2023, Grames & Forister 2024, Li *et al*. 2024). Despite remarkably large sampling effort, many studies have more covariates (measures of climate or the genome) to make their predictions than they do observations. This poses a profound challenge for predicting organismal responses to climate beyond the settings in which they were studied. While some subdisciplines would suggest a priori limiting the number of potential predictor parameters to substantially fewer than available samples (i.e., degrees-of-freedom spending; Giudice *et al*. 2012, Harrell 2015), or the use of model averaging (Dormann *et al*. 2018), here we investigate methods that allow models to be confronted with all available data.

Limited generalizability is common in parameter-rich models and results from overfitting, or the tendency for flexible models to fit too closely to the observed data (see Glossary of key terms). Overfitting occurs when idiosyncratic variation in the observed data is taken as causal associations rather than spurious associations that can arise in finite samples (Hastie *et al*. 2015). Thus, the availability of big data with potentially many more covariates (*P* ; i.e., high-dimensional) than observations (*N*) may counter-intuitively lead to models with poor predictive performance outside the scope of the sample (“the curse of dimensionality”, Altman & Krzywinski 2018). Indeed, ecological studies have experienced a general decline in predictive power over the past century despite concomitant increases in statistical complexity (Low-Décarie *et al*. 2014). More broadly, we face the problem of how to realistically and intelligently constrain the flexibility of our models to capture potential general patterns while learning about genuine context-dependent effects (Weiss 2008). Given that there are perhaps as many ways to analyze a given dataset as there are biologists to analyze it (Gould *et al*. 2025), and that this can affect the magnitude of effect sizes, if not the direction, it is imperative that we grow our understanding of how the models we fit can be expected to perform when applied to covariate-rich data sets.

We have made great strides in ecology and evolution in constructing highly predictive models to address the proliferation of large data sets. Machine learning complements standard methods for statistical modeling that make *a priori* choices of which variables to include based on conceptual, process-based understanding. More generally, advances in machine learning have prompted a reevaluation of how we value models and what it means for a model to ‘understand’ something about the world (Breiman 2001b, Mitchell & Krakauer 2023). Statistical models can be used to learn which variables are associated with a response (i.e., variable or feature selection), to generate accurate predictions about the sampled population (i.e., in-sample prediction), and to make generalizations about populations, localities, or time-points for which we have no prior information about response variables (i.e., out-of-sample prediction). We value models that can reveal potentially causal variables that are associated with the generative processes leading to variation in the response, while also avoiding shortcut learning that can garner accurate predictions from tangentially related variables (Geirhos *et al*. 2020; see Fourcade *et al*. 2018 for an example with species distribution modeling).

Ecologists and evolutionary biologists would benefit from a direct comparison and evaluation of the prospects of different statistical learning methods (Porwal & Raftery 2022), and from greater clarity about critical issues in model evaluation, including overfitting, the extent to which process variance is recovered in model predictions, and the explanatory value of important variables. Computational methods for sparse modeling might be particularly valuable approaches: these methods assume that most variables have no causal relationship with the response and therefore only generate estimates for a subset of variables (Hastie *et al*. 2015). The hope is that selected variables correspond to the key process variables that are causally linked to variation in the response, which should limit overfitting and improve predictive performance when generalizing to unsampled or future observations. This can be contrasted to explicit causal modeling using directed acyclic graphs (DAGs; Laubach *et al*. 2021), or variable selection using information criteria to maximize in-sample prediction and parsimony (see Addicott *et al*. 2022, Arif & MacNeil 2022, for more extensive discussion of prediction at the cost of causal understanding). It is an open question to what extent sparse models can maximize predictive performance and yield interpretable model outputs, particularly for high-dimensional data where the number of covariates (*P*) is much greater than the number of samples (*N*).

We compared the relative performance of several modeling methods by applying them to the same data sets with known, simulated causal relationships, of the type commonly encountered in ecology and evolutionary biology. Our 36 core simulation scenarios (100 simulated replicates each) differed in the number of observations (*N* = 50, 150, or 500), the number of covariates (*P* = 100, 1,000, 10,000, or 100,000; of which 10 were causal and directly influenced the response variable), and the effect size of causal variables (***β***_causal_ = 0.1, 0.3, or 0.8; Table S1). Our statistical learning methods included penalized regression methods based on maximum likelihood (Ridge, Elastic Net, and LASSO) and Bayesian estimation (Bayesian LASSO [BLASSO], Horseshoe, Spike-and-slab, Sum of Single Effects [SuSiE], and Bayesian Sparse Linear Mixed Model [BSLMM]), and one commonly used machine learning method (Random Forest). When selecting methods for inclusion in this study, the primary deciding factor was whether we were potentially excited to use that method for our own research. As such, the methods in this study are biased towards those that are more commonly used in our respective fields (e.g. ecology, genomics, hydrology) or those that have specific features relevant to the data we typically generate. When evaluating the strengths and weaknesses of different methods, we considered prediction (the accuracy of prediction of the response variable given the covariates, both for in-sample, training data and out-of-sample, test data) and inference (i.e., learning which variables are potentially causally associated with variation in the response). While prediction and inference should be treated as complementary goals in statistical analyses (Breiman 2001b), with a continuum between them (Dormann *et al*. 2012), it is worth noting that prediction and inference are not always associated (Arif & MacNeil 2022), even if there is an expectation that strong inference would follow from accurate predictions (Wang *et al*. 2020b).

## 2 Materials and Methods

### 2.1 Description of simulated data

When selecting the attributes of our simulations, we chose a range of variation in *N* and *P* that resemble current ecological and evolutionary datasets (e.g., months of micrometeorological sensor measurements, genome-wide estimates of genetic polymorphisms), though we acknowledge that data dimensionality is continually expanding for most fields. The simulations included 36 scenarios that considered three main factors in a fully crossed design: the number of observations (*N* = 50, 150, or 500), the number of variables or features (*P* = 100, 1,000, 10,000, or 100,000), and the effect size of the ten causal variables (***β***_causal_ = 0.1, 0.3, or 0.8; Table S1). To evaluate the potential benefits of even larger *N*, we simulated two additional scenarios in which *N* was 1,000 or 10,000, *P* was 1,000, and ***β***_causal_ was 0.3. To thoroughly incorporate and evaluate variable outcomes among simulations, we obtained 100 replicate data sets for all scenarios. Each replicate data set consisted of *N* observations for training (i.e., variable selection and in-sample prediction) and an additional 500 observations for testing out-of-sample prediction. We did not explore how variation in the size of the out-of-sample test data influenced model performance, but this would be worthy of exploration in future studies.

For each replicate, we first created an observation (*N* + 500)*×* variable (*P*) matrix ***X*** consisting of *P* /50 clusters of correlated variables (50 per cluster). Each cluster of variables was generated by taking *N* + 500 draws from a multivariate normal distribution with mean vector ***µ*** = 0 and covariance matrix **Σ**. We generated covariance matrices using a spherical parameterization (Pinheiro & Bates 1996), which transforms a *P* (*P* +1)/2-dimension vector of unconstrained parameters ***θ*** into a positive semi-definite covariance matrix **Σ**. The goal of this approach was to create clusters of variables with a range of correlation strengths, from strongly negatively to strongly positively correlated, a situation that is common in biological relationships and that presents a challenge for many modeling approaches (but see Dormann *et al*. 2013, for solutions to limit colinearity in anal-yses). We found that drawing values of ***θ*** from a uniform distribution between -1 and 1 produced sets of variables with a range of correlation strengths. After generating clusters of variables, we concatenated them to create the variable matrix ***X*** and centered and scaled (mean = 0; sd = 1) the columns of variables.

Next, we sampled a *P* -dimension vector of coefficients ***β*** representing the causal effects of the variables on response variable ***y***. We randomly selected 10 variables out of *P* to have a non-zero coefficient of ***β***_causal_. The remaining values of ***β*** were set to zero. The response variable ***y*** was a linear, additive function of the product of the ***β*** coefficients and the *P* variables, plus error of ***E***, drawn from a standard normal distribution for each individual: ***y*** = ***Xβ*** + ***E***. For each data set, the reducible error was calculated as the proportion of variance in the response explained by a linear model using only the 10 causal variables.

We made several decisions in simulating data that could influence our results and interpretation. For example, causal parameters in the simulated data sets were specified as simple linear effects, as opposed to non-linear or threshold effects that could be more or less difficult to identify for some methods. However, while linear approximations of non-linear processes introduce bias, they can often outperform more flexible non-linear or non-parametric approaches that introduce more variance, particularly for high-dimensional data (i.e., “the bias-variance trade-off”; James *et al*. 2021). Furthermore, we intentionally avoided the complexities of causal inference in the presence of confounding variables and interactions. Instead, we studied a simplified system in which sparse effects could estimate causal effects; this can be thought of as a best case scenario for the purpose of statistical learning. Finally, we have explored a fairly simple range of data attributes that might be encountered in the life sciences, and acknowledge that the consideration of other axes of variation (e.g. thresholds, degree of colinearity, non-continuous distributions) will undoubtedly lead to new insights about how we can use modeling approaches to better understand the world.

### 2.2 Analyses

Each simulated data set was analyzed using nine different methods. Eight of these are penalized regression methods using standard likelihood (LASSO, Tibshirani 1996; Ridge, Hoerl & Kennard 1970; Elastic Net, Zou & Hastie 2005) or Bayesian estimation (Bayesian LASSO [BLASSO], Park & Casella 2008; Horseshoe, Carvalho *et al*. 2010; Spike-and-slab, Ishwaran & Rao 2005; Bayesian sparse linear mixed model [BSLMM], Zhou *et al*. 2013; sum of single effects [SuSiE], Wang *et al*. 2020a). The final method, Random Forest (Breiman 2001a), served as a benchmark to compare other methods to and is a commonly used, highly flexible machine learning approach based on an ensemble of decision trees. All analyses were conducted in R v4.2.2 (R Core Team 2023). Each data set was provided to the methods using the Nextflow v22.10.4.5836 workflow description language (Di Tommaso *et al*. 2017) to distribute the work and aggregate the output in a computing cluster using SLURM (Yoo *et al*. 2003). We used implementations of Elastic Net, LASSO, and Ridge in the glmnet v4.1-6 package (Friedman *et al*. 2010), of BLASSO, Horseshoe, and alternatives of LASSO and Ridge in the monomvn v1.9-17 package (Gramacy 2023), of Spike-and-slab in the spikeslab v1.1.6 package (Ishwaran *et al*. 2010), of SuSiE in the susieR v0.12.27 package (Wang *et al*. 2020a), of Random Forest in the randomForest v4.7-1.1 package (Liaw & Wiener 2002), and of BSLMM in the software gemma v0.98.6 (Zhou *et al*. 2013). We used ‘off-the-shelf’, default settings for all analyses (as in Porwal & Raftery 2022). This means that we did not perform hyperparameter optimization, which could potentially lead to biases when comparing methods, as the importance of optimization varies across models (Bischl *et al*. 2023). BLASSO and Horseshoe were not performed for the large *N* scenarios (*N* = 1,000 or 10,000) due to extremely long run times. All R scripts for creating the simulated data, running analyses, generating summary statistics, and making figures, as well as the Nextflow wrappers and config files are archived at Zenodo (doi:10.5281/zenodo.19076540).

To evaluate each model’s potential utility for parameter estimation, variable selection, and prediction, we calculated several complementary summary statistics that were largely applicable across all of the methods. Metrics for BLASSO and Horseshoe were calculated two ways: model-averaged (ma) estimates are based on all samples from the reversible jump MCMC, whereas non-zero (nz) estimates use only samples in which the variable and associated coefficient were included in the model. Parameter estimation was evaluated based on the root mean square error (RMSE) between estimated and actual parameter values (***β***) for all analyses except Random Forest, which reports importance measures instead of estimates. Variable selection was first assessed for methods that can return true zeros for parameter estimates (BSLMM, Elastic Net, LASSO, Spike-and-slab) or importance measures (Random Forest). Variables were assigned as positives (*/*= 0) or negatives (= 0), and these classifications were used to calculate true positive rates (TPR; i.e., sensitivity), true negative rates (TNR; i.e., specificity), and F_1_, which is the harmonic mean of precision (i.e., the fraction of selected variables that are truly causal) and sensitivity: 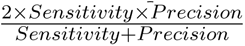. It is important to note that small values of F_1_ (i.e., poor variable selection) can occur due to low TPR, low TNR, or both. Variable selection was also assessed based on posterior inclusion probabilities (PIPs) for four Bayesian methods (BLASSO, BSLMM, Horseshoe, SuSiE) using one example data set (scenario 24, replicate 1). A series of minimum PIP thresholds (i.e., variables with PIP *≥* threshold are scored as positives) were evaluated to characterize potential effects on resulting F_1_ values. In-sample and out-of-sample prediction was quantified using R^2^ between the actual and predicted values of the response variable. In-sample prediction was based on the *N* observations used to train the model, whereas out-of-sample prediction was based on a separate set of 500 observations. Finally, we recorded the runtime required to fit each model to each data set.

## 3 Results

A perfect model would: 1) identify only the ten truly causal variables and accurately estimate their effect sizes; 2) accurately attribute the variation in the response that arises directly from the causal variables (i.e., reducible error); and 3) disregard variation in the response arising from other unmeasured or stochastic processes (i.e., irreducible error; James *et al*. 2021). Across all simulations, the magnitude of reducible error was overwhelmingly associated with the effect size of causal variables (R^2^ between the known, additive effects of simulated causal variables and the simulated response variable was *≈* 0.10, 0.47, and 0.86 with ***β***_causal_ = 0.1, 0.3, and 0.8, respectively; Fig. S1). Reducible error was more variable among replicates when *N* or *P* were low.

The different methods varied greatly in their performance for variable selection and prediction. For one example data set (*N* = 150, *P* = 10,000, ***β***_causal_ = 0.8; see Fig. 1), LASSO monomvn had the greatest success at delineating between causal and non-causal variables (true positive rate [TPR] = 0.9; true negative rate [TNR] = 0.997). While Random Forest correctly identified all ten causal variables (TPR = 1), it implicated a large proportion of non-causal variables as being associated with the response (TNR = 0.118). In contrast, BSLMM was successful at excluding non-causal variables (TNR = 0.965), but could only identify six of the causal variables (TPR = 0.6). For prediction, the true reducible error for the example data set was R^2^ = 0.832, which served as the target for both in-sample and out-of-sample prediction (based on 500 observations not used to train the model). For LASSO monomvn, in-sample prediction was very close to the reducible error (R^2^ = 0.826), which translated to the highest success for out-of-sample prediction (R^2^ = 0.754). In-sample prediction exceeded the reducible error for BSLMM (R^2^ = 0.970), and this overfitting led to reduced out-of-sample prediction (R^2^ = 0.617) relative to LASSO monomvn. Random Forest suffered from poor predictive performance, with both in-sample (R^2^ = 0.124) and out-of-sample (R^2^ = 0.332) comparisons, falling far short of the reducible error (see discussion of underfitting below). Overall, LASSO monomvn provided the best balance between variable selection and prediction for the example data set.

**Figure 1:**
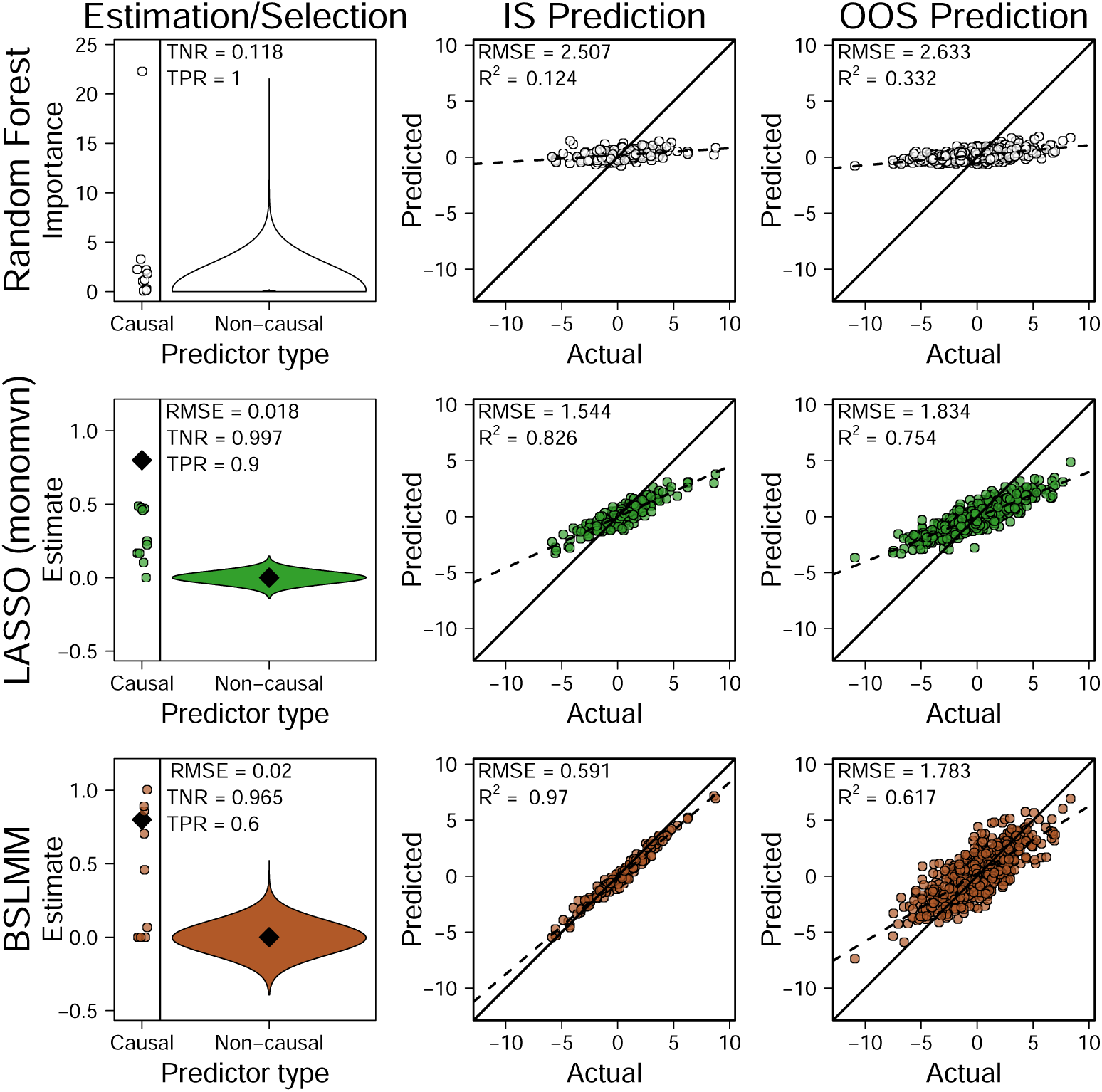
Performance varies greatly across three methods for parameter estimation, variable selection, in-sample (IS) prediction, and out-of-sample (OOS) prediction. Results are shown for the first replicate of scenario 24, which had 10 causal variables (***β*** = 0.8) and 9,990 non-causal variables (***β*** = 0). The distributions of causal and non-causal importance values are shown for Random Forest, whereas the distributions of causal and non-causal parameter estimates are shown for LASSO monomvn and BSLMM (black diamonds signify the true effect sizes). In-sample prediction was based on 150 observations used to train the model, and out-of-sample prediction was based on a separate 500 observations (the maximum reducible error for this scenario was R^2^ = 0.832). RMSE: root mean square error; TNR: true negative rate (i.e., specificity); TPR: true positive rate (i.e., sensitivity)

Overfitting was rampant across all scenarios, as evidenced by large in-sample R^2^ and low out-of-sample R^2^ (Fig. 2A,B). It was also common for models to recover only a fraction of the reducible error in out-of-sample prediction, particularly for simulations with larger *P* (Fig. 2B). The accuracy of in-sample and out-of-sample prediction converged towards the reducible error target for simulations with larger ***β***_causal_ and *N* and smaller *P* (Fig. 3). Out-of-sample predictive performance was not necessarily associated with more accurate variable selection, as out-of-sample R^2^ matched the reducible error even with low F_1_ for some scenarios (Fig. S2). Variable selection was first assessed for methods that return truly sparse parameter estimates (i.e., ***β*** = 0; BSLMM, Elastic Net, LASSO, Spike-and-slab) or importance values (Random Forest), and was generally poor except for when ***β***_causal_ and *N* were high and *P* was low; Fig. 2C). When ***β***_causal_ was low, a negative relationship between TPR and TNR emerged across methods, suggesting an intuitive trade-off between identifying causal variables and excluding non-causal variables (Fig. 4). Variable selection was also assessed based on posterior inclusion probabilities (PIPs) for four Bayesian methods (BLASSO, BSLMM, Horseshoe, SuSiE) using the example data set from Fig. 1. The use of a small PIP threshold of 0.05 (i.e., only variables with PIP *≥* 0.05 are scored as positives) improved variable selection for BSLMM and SuSiE, whereas larger thresholds were needed to recover more limited gains for BLASSO and Horseshoe (Fig. S3). Parameter estimation was remarkably consistent across different analyses, and was instead most strongly influenced by data dimensionality: estimation was worse with greater ***β***_causal_ and lower *N* and *P* (Fig. 2D). This pattern arose because most methods are worse at estimating variables with ***β*** */*= 0 than those with ***β*** = 0, resulting in larger root mean square error (RMSE) when the proportion of causal to non-causal variables was relatively large (i.e., when *P* was small) or when the effect size of causal variables was very different from zero (i.e., when ***β***_causal_ was large). The ratio of causal to non-causal variables will be particularly influential on parameter estimation for methods using penalization (e.g. Ridge, LASSO, Elastic Net, BLASSO), as the penalty will depend on the sample size, with analyses of smaller sample sizes having more parameters shrunk to 0. Analysis of the 3,600 data sets completed in 2.49 CPU-years, with BLASSO and Horseshoe contributing 46.6% and 46.7% of the total run-time, respectively (Fig. 2E).

**Figure 2:**
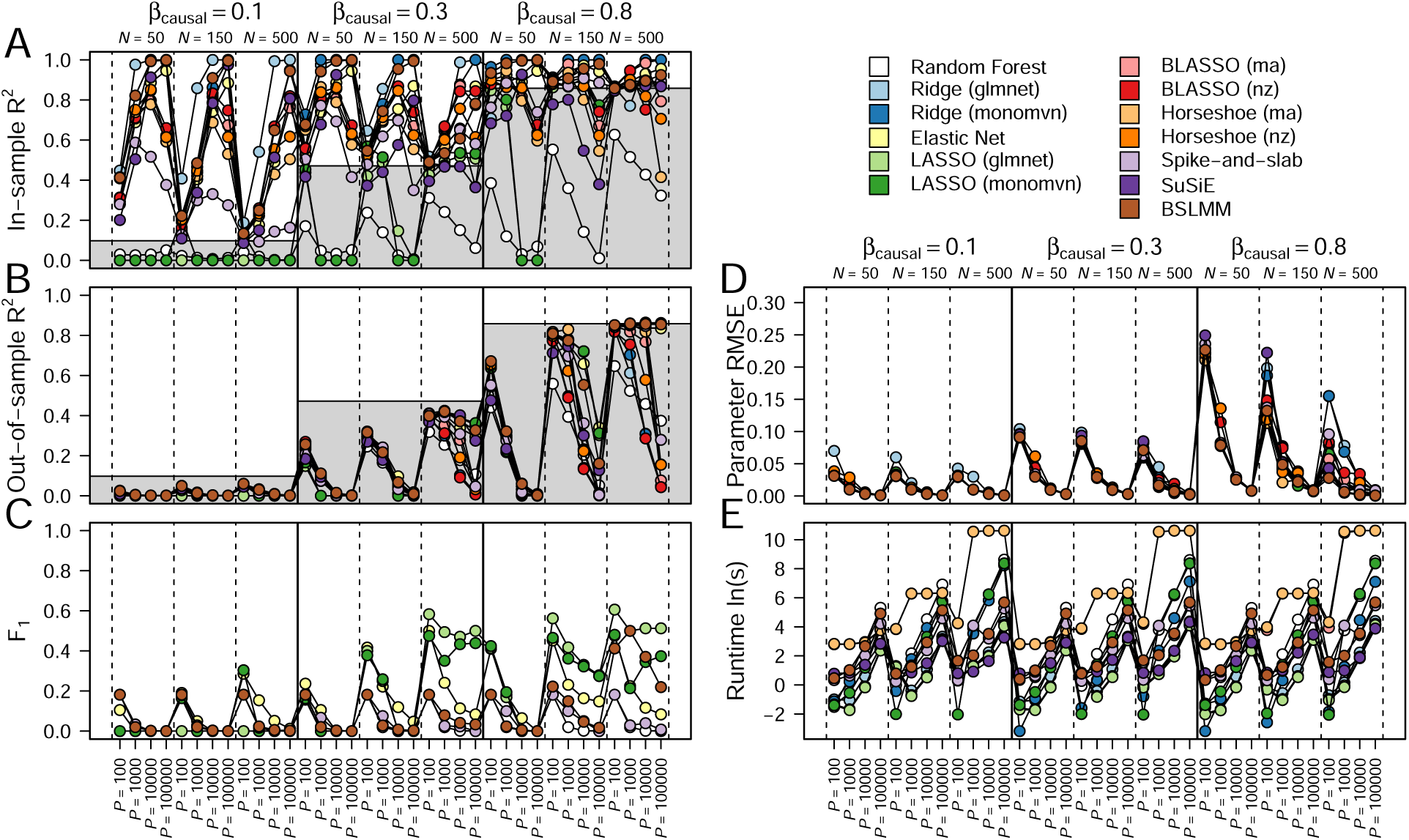
An overview of model performance for the 36 core scenarios. Nine methods were con-sidered, as well as two different implementations of Ridge and LASSO in the glmnet and monomvn packages in R. Metrics for BLASSO and Horseshoe were calculated two ways: model-averaged (ma) estimates are based on all samples from the reversible jump MCMC, whereas non-zero (nz) estimates use only samples in which the variable and associated coefficient were included in the model. (A) In-sample and (B) out-of-sample prediction were evaluated with R^2^ between the actual and predicted values of the response. In these panels, the grey and white regions represent the mean reducible and irreducible error, respectively, across all scenarios within a ***β***_causal_ level. While reducible error represents the expected maximum value for out-of-sample prediction, in-sample prediction can exceed the reducible error when too flexible models are employed (i.e., overfitting). This means that the target for prediction is to recover a model with in-sample and out-of-sample R^2^ equal to the maximum reducible error. (C) Variable selection was evaluated using F_1_, which is the harmonic mean of precision (i.e., the fraction of selected variables that are truly causal) and sensitivity (i.e., true positive rate): 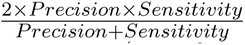. F_1_ was only calculated for analyses that can return truly sparse parameter estimates (i.e., ***β*** = 0; BSLMM, Elastic Net, LASSO, Spike-and-slab) or importance values (Random Forest). (D) Parameter estimation was evaluated for all methods except Random Forest using the root mean square error (RMSE) between estimated and actual parameter values. (E) Model speed was evaluated based on the natural log of runtime in seconds. Each circle represents the median value from 100 replicate simulations.

**Figure 3:**
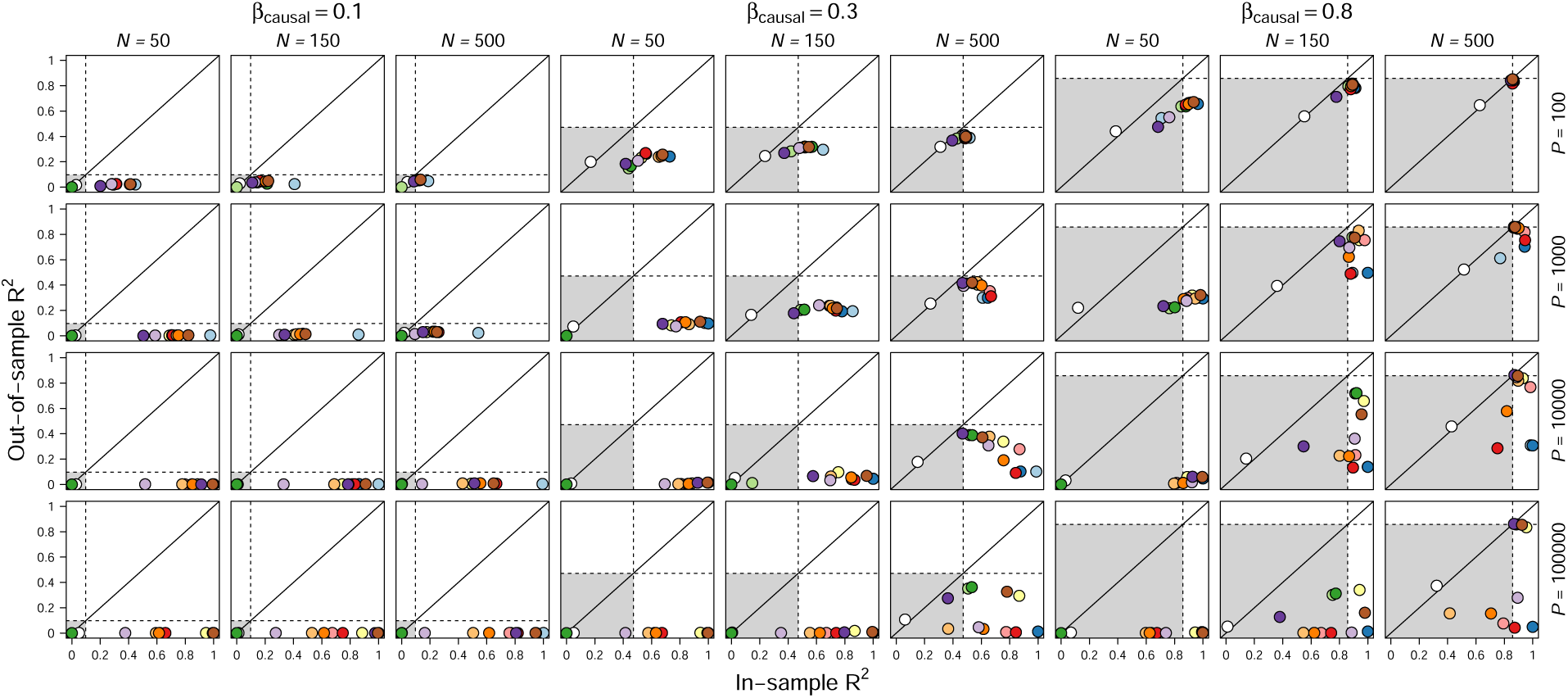
The extent of overfitting and the fraction of reducible error recovered differs dramatically among methods and data attributes. The grey and white regions represent the mean reducible and irreducible error, respectively, across all scenarios within a ***β***_causal_ level. While reducible error represents the expected maximum value for out-of-sample prediction, in-sample prediction R^2^ can exceed the reducible error when too sensitive models are employed (i.e., overfitting). This means that the target for prediction is to recover a model with in-sample and out-of-sample R^2^ equal to the maximum reducible error, as was the case for many of the methods in the upper right hand panel (***β***_causal_ = 0.8; *N* = 500; *P* = 100). Each circle represents the median value from 100 replicate simulations. See Fig. 2 for color legend.

**Figure 4:**
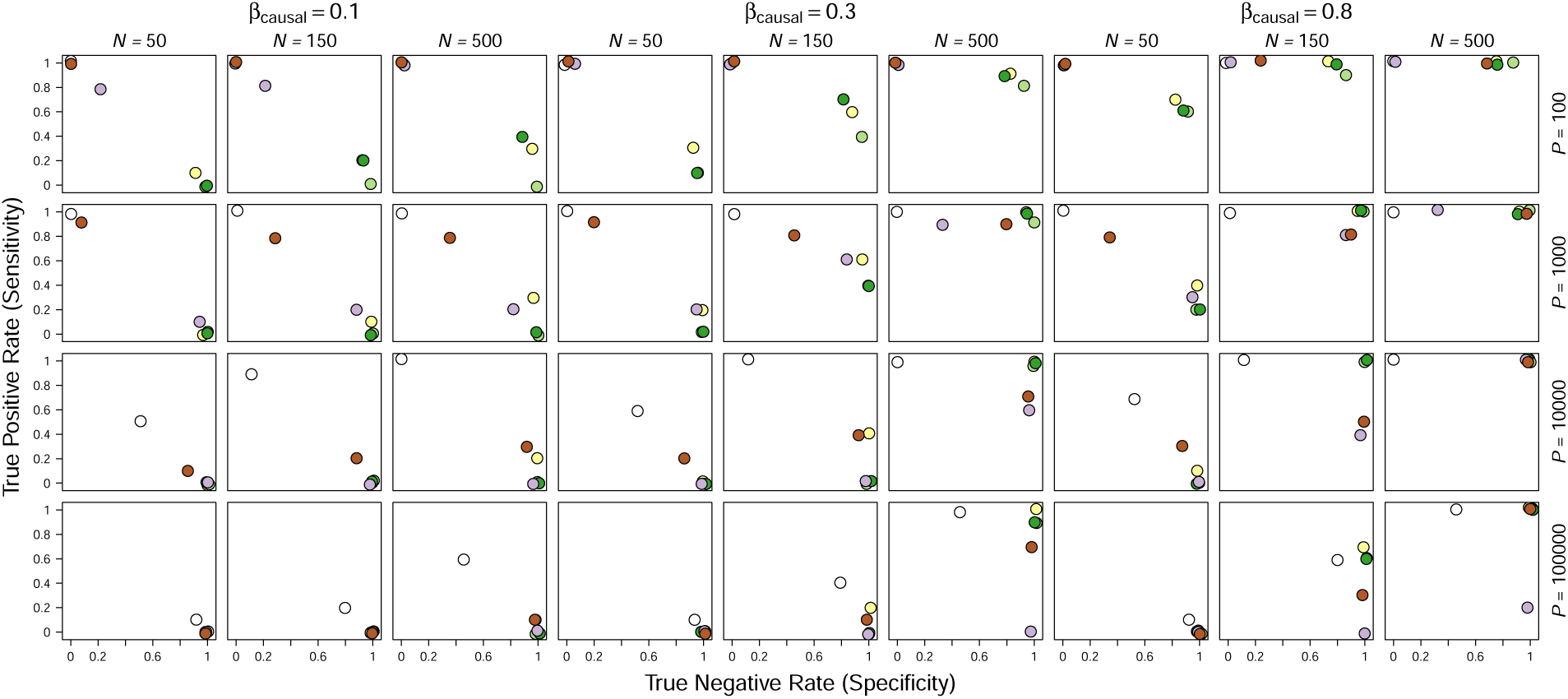
Variable selection performance varied greatly across scenarios, and was only possible for some methods when there were many observations (*N*), few variables (*P*), and effect sizes (***β***_causal_) were large. A negative correlation between true positive rate and true negative rate emerged for many simulations, especially when ***β***_causal_ was small, indicative of a trade-off between identifying causal variables (sensitivity) and excluding non-causal variables (specificity). This trade-off disappears when conditions are more favorable for variable selection: when ***β***_causal_ and *N* are large and when *P* is small. Variable selection was only evaluated for analyses that can return truly sparse parameter estimates (i.e., ***β*** = 0; BSLMM, Elastic Net, LASSO, Spike-and-slab) or importance values (Random Forest). Each circle represents the jittered median value from 100 replicate simulations. See Fig. 2 for color legend.

## 4 Discussion

High-throughput and automated data acquisition promises to yield valuable information about processes that generate variation. This promise is diminished in the common situation in ecology and evolutionary biology when sampling is of few individuals (*N*) and many potential covariates (*P* ; e.g., genomic polymorphisms at 10^6^ sites, months of micrometeorological sensor measurements at 10Hz). Our simulations highlight that the most consistent way to obtain highly predictive and explanatory models is to maximize the number of independent observations. We recognize that to tell ecologists and evolutionary biologists ‘just collect more data’ is not always a helpful statement. However, when we extended our simulations to sample sizes of 1,000 or 10,000 observations, in-sample and out-of-sample R^2^ converged to the maximum reducible error, and variable selection improved for most analyses (Fig. 5). This positive relationship between model performance and sample size is likely more nuanced than is shown in Fig. 5, as the limited number of sample sizes (*N*) found in our simulations could obscure potential non-linear or algorithm specific patterns that should be investigated in future studies. While sparse modeling techniques allow the fitting of models in settings with more covariates than observations (*P > N*), they cannot rescue analyses based on small sample sizes, especially when *P* is large or when effect sizes are small relative to background levels of stochastic variation (Fig. 2). This means that for many typical analyses in ecology and evolutionary biology, variable selection will suffer from low precision and sensitivity, and prediction models will be overfit and have poor generalizability. In cases where sparse methods struggle with a low signal-to-noise ratio, other methods will also struggle (“the bet-on-sparsity principle”; Hastie *et al*. 2009), though model averaging frameworks could benefit prediction when model covariance is low and the ratio of bias to variance is small (Dormann *et al*. 2018). We do not want to imply that statistics should not be done when sample sizes are small. Indeed, in some cases, such as when species are rare or endangered, small samples are better than nothing. However, we counsel following the precautionary principle when applying models from small training sets to out of sample data, as prediction is expected to be poor.

**Figure 5:**
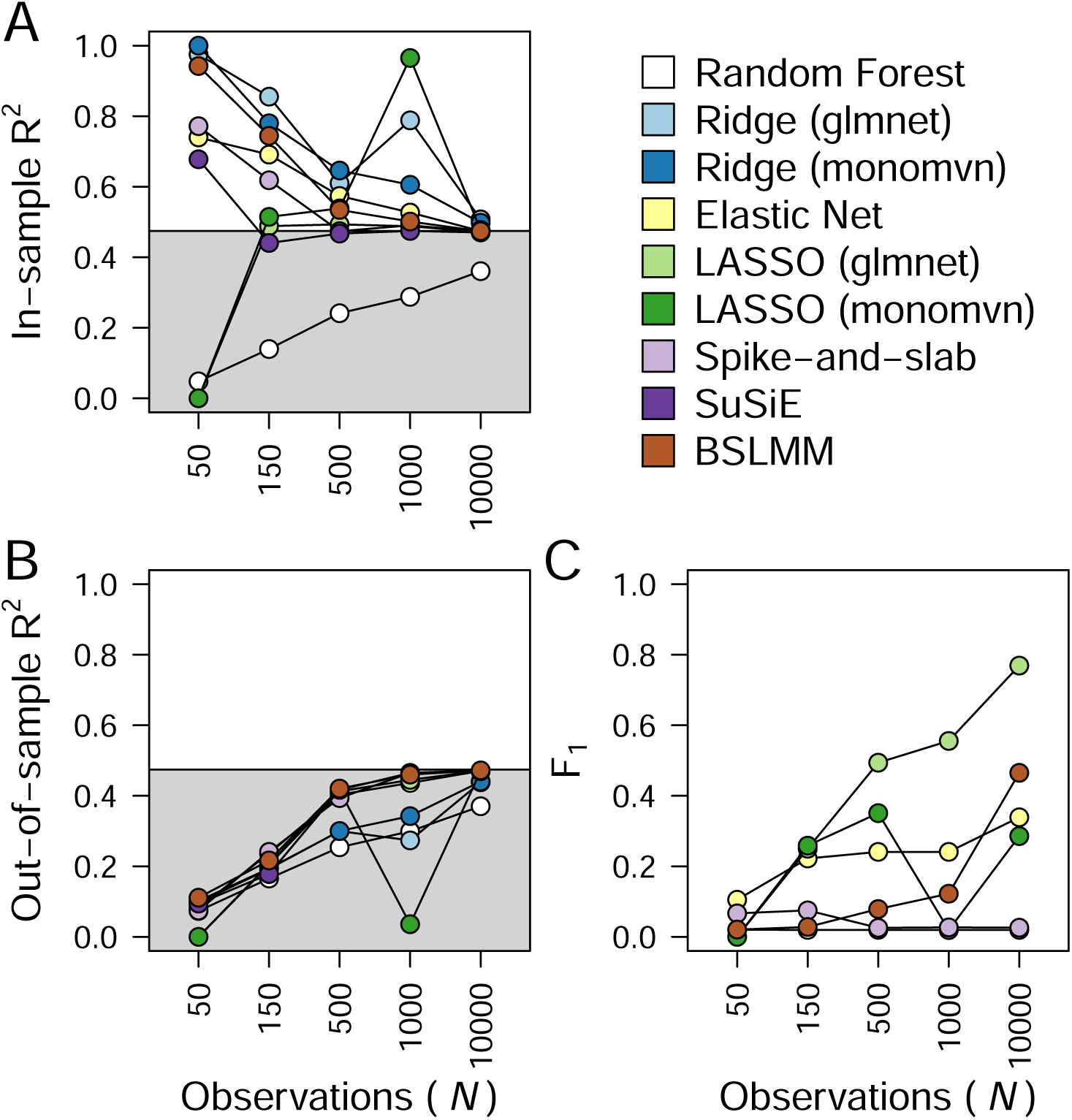
Model performance for (A) in-sample prediction, (B) out-of-sample prediction, and (C) variable selection improves with increasing observations (*N*) for most models (BLASSO and Horse-shoe were not considered because of long runtimes). The five scenarios shown here all had *P* = 1,000 and ***β*** = 0.3. In-sample and out-of-sample prediction were evaluated with R^2^ between the ac-tual and predicted values of the response. In panels A and B, the grey and white regions represent the mean reducible and irreducible error, respectively, across all five scenarios. While reducible error represents the expected maximum value for out-of-sample prediction, in-sample prediction can exceed the reducible error when too flexible models are employed (i.e., overfitting). This means that the target for prediction is to recover a model with in-sample and out-of-sample R^2^ equal to the maximum reducible error. Variable selection was evaluated using F_1_, which is the harmonic mean of precision (i.e., the fraction of selected variables that are truly causal) and sensitivity (i.e., true positive rate): 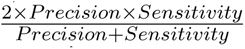. F_1_ was only calculated for analyses that can return truly sparse parameter estimates (i.e., ***β*** = 0; BSLMM, Elastic Net, LASSO, Spike-and-slab) or importance values (Random Forest). Each circle represents the median value from 100 replicate simulations.

It is perhaps näive to use statistical learning for prediction without large training sets, particularly when causal effect sizes are small relative to variance from extraneous sources. The temptation to do so might stem from working with big data (*N × P*), but not appreciating that all statistical approaches are expected to yield relatively poor out-of-sample prediction when *N* is small (e.g., *<* 500) and effect sizes are modest. Some of the most remarkable models in society, such as those for large language modeling (Zhao *et al*. 2023), natural voice recognition (Xiong *et al*. 2016), image segmentation (Kirillov *et al*. 2023), and board game algorithms (Silver *et al*. 2018), are typically trained on enormous sample sizes. For example, Tabak *et al*. (2019) trained a convolutional neural network with more than 3 million images to achieve more than 80% out-of-sample accuracy when detecting ungulates from camera trap imagery. We do believe there is a place for sparse methods in the life sciences when many observations (*N*) can be obtained (Fig. 5). Our simulations provide context for evaluating different dimensions of model quality and the comparison of model approaches suggests which methods will be most useful and when.

For most predictive contexts, the primary objective is to account for the reducible error in the data, as this is the variation in the response associated with generative processes (James *et al*. 2021). We were able to directly quantify the reducible error in our simulated data sets and easily identify cases of overfitting in which in-sample R^2^ *>* reducible error R^2^ (Figs. 2A & 3). With empirical data, the true reducible error and prediction errors arising from model variance and bias will be unknown (James *et al*. 2021). While these can be estimated via cross validation or bootstrapping (Fieberg & Johnson 2015, Harrell 2015), overfitting may be evident when in-sample R^2^ exceeds out-of-sample R^2^. It is worth emphasizing that in our simulations and analyses, we minimized the potential for model bias and underfitting by simulating data from simple additive generative processes that are mirrored in the statistical learning methods we used. In other words, we simulated the best case scenarios for explaining reducible error, and we still typically fell short.

To minimize errors in prediction, we can strive for large sample sizes of representative data for model training (i.e., homogeneous with the out-of-sample, test data). Additionally, on average, in-sample prediction accuracy cannot be less than out-of-sample prediction accuracy, and both will converge on the true reducible error with increasing ***β***_causal_ and *N* and decreasing *P* (Fig. 3). Recovery of similar in-sample and out-of-sample R^2^ is consistent with minimal overfitting, but could arise from model bias (e.g., choosing too simple of a model that has large bias but small variance; Lever *et al*. 2016, James *et al*. 2021). Similar out-of-sample R^2^ from multiple, genuinely different analysis methods would be consistent with having minimized prediction error given the information in the available data, but could still derive from underfit, biased models that account for only a fraction of the true reducible error (see results from Random Forest in the example data set; Fig. 1). This is consistent with the observation by Dietterich (1995) that optimizing the fit to the training data will limit the predictive ability of a decision tree. It is worth noting that Random Forest always yielded in-sample R^2^ roughly equal to out-of-sample R^2^ (Figs. 2A,B & 3), as this is the only method that uses cross-validation as a default, thus limiting our ability to fairly compare results across methods. The distinction between prediction errors that arise in-sample and out-of-sample (Fig. 3), and the strong potential for overfitting, call into question model choice decisions that are very commonly made based on in-sample data alone without any cross-validation procedure (i.e., potentially choosing the most overfit model with little deference for out-of-sample prediction; see Fig. 3; Tredennick *et al*. 2021). Importantly, while cross-validation is critical for safeguarding against misleading model results, reduced *R*^2^ values from cross-validated models could result in studies being less likely to be published or going into a lower profile journal (i.e., the “file drawer problem”; Csada *et al*. 1996, Low-Décarie *et al*. 2014), suggesting the need for a shift in how researchers evaluate prediction results in the context of cross-validation.

The conditions that allow for strong prediction, namely when ***β***_causal_ and *N* are large and *P* is small, are the same in which variable selection is possible (Figs. 2C & 3), though reliable out-of-sample prediction did not necessarily depend on perfect variable selection (i.e., including all causal and excluding all non-causal variables; Fig. S2). A variable selection trade-off emerged for the data sets in which variable selection was most difficult (e.g., ***β***_causal_ = 0.1), as evidenced by a negative relationship between true positive and true negative rates (Fig. 4). This result has broad implications because effect sizes are expected to be small and diffuse for many biological systems (e.g., in genetics, Boyle *et al*. 2017). Moreover, the consequences of different variable selection errors will have disparate repercussions in exploratory versus diagnostic settings, so researchers will need to weigh the costs and benefits of either identifying all causal variables at the expense of including some false positives (e.g., when developing candidate variables for further study) or missing some causal variables to ensure the absence of any false positives (e.g., when identifying biomarkers for disease detection). In other words, best statistical practices, including when using sparse models, rely on identifying whether an analysis is hypothesis generating or hypothesis testing prior to implementation (Tredennick *et al*. 2021). For the Bayesian methods that generate posterior inclusion probabilities (PIPs), the threshold for deciding whether or not to include a variable may vary across disciplines and fields. In evolutionary genetics, researchers may choose to only consider genetic loci that have a PIP *>* 0.1 (Lucas *et al*. 2018, McFarlane & Pemberton 2021), and this simple choice would have substantially improved variable selection for BSLMM and SuSiE, but not BLASSO or Horseshoe, for one example scenario (Fig. S3). Overall, accurate variable selection requires large numbers of observations (Fig. 5), perhaps even more so than prediction, as has been found previously in trait mapping and phenotypic prediction (Wray *et al*. 2013, Gompert *et al*. 2017).

One striking feature of our results was the absence of a single method that excelled at all modeling purposes, consistent with the “no free lunch theorem” for supervised learning (Wolpert 1996, Wolpert & Macready 1997). Trade-offs in model building have long been recognized (Levins 1966, James *et al*. 2021, Tredennick *et al*. 2021) and serve as an important reminder for researchers to wield methods that align with their research objectives. Consequently, it can be useful to simulate data and measure the correlation (and other measures of the relationship) of the response variable with process parameters (***β***_causal_) under relevant sample sizes, so as to gauge information about the expected reducible error. It may be the case that researchers will need to employ multiple, complementary statistical learning methods for questions involving both prediction and variable selection. A combined approach to model building could be particularly valuable, whether that be through model averaging (Dormann *et al*. 2018), hybrid modeling structures that combine process-based and empirical approaches (Reichstein *et al*. 2019), or a more iterative approach: for example, using a sparse method to identify a subset of candidate variables and following up with a more flexible method such as Random Forest for prediction. We emphasize that while the use of sparse methods cannot resolve logistical challenges surrounding data collection in ecology and evolutionary biology (there will always be data sets where *P* is much greater than *N*, or where *N* is inherently small), the uptake of these methods is a path forward that can contribute to high-quality inference, explanatory models that capture key elements of data generating processes, and prediction with minimal error. Finally, we acknowledge that while many of our key findings recapitulate concepts that are already well known by many statisticians (James *et al*. 2021), they are less widely appreciated in ecology and evolutionary biology, and we hope that we have raised awareness of the promise and limitations of sparse modeling tools for statistical learning.

## 5 Glossary

- **Bias-variance tradeoff:** model fitting can be highly flexible and responsive (the variance part) or insensitive (the bias part) to data that are used for statistical learning. These two attributes are inversely related to one another and exhibit a trade-off (James *et al*. 2021).
- **Causal relationship:** one process or measure affects another, directing or influencing a response. This is in contrast to associations between measures that arise from sources other than causality (spurious or confounded associations, etc.).
- **Covariates (***P* ): measures of interest that can be used as potential explanatory variables for a response variable (e.g., many climate variables as covariates in predicting survivorship).
- **Cross-validation:** a scheme by which fractions of the data are iteratively left out during the training (fitting) stage of model-building. The left out data are used as a gauge to avoid having the model fit the training data excessively well, while having a poor fit to left out data (i.e., guard against over-fitting).
- **Generalizability:** a goal in science to obtain explanations and models that apply to broader contexts beyond a narrow scope of an individual sampling unit, study, population, etc. (please note that other studies instead call this transferability; e.g. Spake *et al*. 2022). While generalizability is an aspiration, whether it is achievable or not is an open question and a scientific result itself.
- **Generative process:** the production of a measurable system feature by biological, chemical, and physical phenomena, which generate or cause the result with some level of determinism.
- **High-dimensional data:** herein we are drawing attention to data sets with many covariates (dimensions), potentially providing sufficient information to predict the response variable, but also raising challenges we describe.
- **In-sample or out-of-sample prediction:** model fit can be quantified using the same samples that were used for model-fitting as is done commonly in statistics (in-sample; using *R*^2^, residuals, information criteria like AIC, etc.), or a held-out set of samples (out-of-sample).
- **Inference:** an understanding based on evidence, reasoning, or both; in this context, an understanding of relationships based on the association between the parameters and the response variable of interest.
- **Irreducible error:** excess variability in predictions beyond what can be accounted for by causal processes (James *et al*. 2021).
- **Machine learning:** statistical algorithms that use large sets of data to recognize patterns, adjust models, and make predictions of additional, out-of-sample data.
- **Overfitting:** a phenomenon in which a model places too much weight on variables that are not related to causal processes, thus limiting the model’s potential for out-of-sample prediction (i.e. poor generalizability; James *et al*. 2021).
- **Precision:** the fraction of selected variables that are truly causal.
- **Reducible error:** variation in the response variable that can be truly attributed to causal parameters, in contrast to variation that arises due to extraneous causes (see Irreducible error).
- **Root mean squared error (RMSE):** a commonly used metric for estimating how far response parameter estimates deviate from the true value (larger values indicate more error).
- **Sample size (***N* ): the number of units of observation that are available to estimate numeric relationships between the response variable and covariates in support of inference and prediction.
- **Sensitivity:** the fraction of causal parameters that are accurately identified by the model (i.e., the true positive or recall rate).
- **Sparse modeling:** a statistical framework in which a relatively small proportion of covariates is assumed to have a strong relationship with the response variable (Hastie *et al*. 2015).
- **Specificity:** the number of non-causal parameters that are accurately excluded by the model (i.e., the true negative rate).
- **Statistical learning:** understanding data using computational methods (James *et al*. 2021).
- **Variable (or feature) selection:** the identification of a subset of predictors that are associated with variation in the response variable.

## Acknowledgments

All authors were supported by the Modelscape Consortium with funding from the National Science Foundation (OIA-2019528). We thank: members of the Modelscape Consortium for input during early discussions of this project; Chris Moore for fun conversations about the philosophy of model building; Dan Gibson (I miss you, friend), Loren Rieseberg, and two anonymous reviewers for thoughtful feedback on earlier drafts. Analyses were performed using University of Wyoming’s Advanced Research Computing Center and its Beartooth Computing Environment, Intel x86_64 cluster (https://doi.org/10.15786/M2FY47).

## Author contributions

Joshua P. Jahner, C. Alex Buerkle, S. Eryn McFarlane, Andrew Siefert, Matthew L. Forister, Daniel C. Laughlin, Breanna F. Powers, and Isabella A. Oleksy designed research; Andrew Siefert created simulations; Joshua P. Jahner, C. Alex Buerkle, Dustin G. Gannon, Eliza M. Grames, S. Eryn McFarlane, Andrew Siefert, Joshua G. Harrison, Chhaya M. Werner, and Isabella A. Oleksy performed analyses; C. Alex Buerkle, Matthew L. Forister, and Daniel C. Laughlin acquired funding; all authors contributed to writing and revision.

