## Supplementary material for "Interpretable and predictive models based on high-dimensional data in ecology and evolution": Supp

| Scenario | $P$ | $N$ | $\boldsymbol{\beta}_{\text{causal}}$ |
|---|---|---|---|
| Core 1 | 100 | 50 | 0.1 |
| Core 2 | 100 | 50 | 0.3 |
| Core 3 | 100 | 50 | 0.8 |
| Core 4 | 100 | 150 | 0.1 |
| Core 5 | 100 | 150 | 0.3 |
| Core 6 | 100 | 150 | 0.8 |
| Core 7 | 100 | 500 | 0.1 |
| Core 8 | 100 | 500 | 0.3 |
| Core 9 | 100 | 500 | 0.8 |
| Core 10 | 1,000 | 50 | 0.1 |
| Core 11 | 1,000 | 50 | 0.3 |
| Core 12 | 1,000 | 50 | 0.8 |
| Core 13 | 1,000 | 150 | 0.1 |
| Core 14 | 1,000 | 150 | 0.3 |
| Core 15 | 1,000 | 150 | 0.8 |
| Core 16 | 1,000 | 500 | 0.1 |
| Core 17 | 1,000 | 500 | 0.3 |
| Core 18 | 1,000 | 500 | 0.8 |
| Core 19 | 10,000 | 50 | 0.1 |
| Core 20 | 10,000 | 50 | 0.3 |
| Core 21 | 10,000 | 50 | 0.8 |
| Core 22 | 10,000 | 150 | 0.1 |
| Core 23 | 10,000 | 150 | 0.3 |
| Core 24 | 10,000 | 150 | 0.8 |
| Core 25 | 10,000 | 500 | 0.1 |
| Core 26 | 10,000 | 500 | 0.3 |
| Core 27 | 10,000 | 500 | 0.8 |
| Core 28 | 100,000 | 50 | 0.1 |
| Core 29 | 100,000 | 50 | 0.3 |
| Core 30 | 100,000 | 50 | 0.8 |
| Core 31 | 100,000 | 150 | 0.1 |
| Core 32 | 100,000 | 150 | 0.3 |
| Core 33 | 100,000 | 150 | 0.8 |
| Core 34 | 100,000 | 500 | 0.1 |
| Core 35 | 100,000 | 500 | 0.3 |
| Core 36 | 100,000 | 500 | 0.8 |
| Large $N$ 1 | 1,000 | 1,000 | 0.3 |
| Large $N$ 2 | 1,000 | 10,000 | 0.3 |

Table S1: Specifications for the 38 simulation scenarios, including the number of predictors ($P$), the number of observations ($N$), and the effect size of causal predictors ($\boldsymbol{\beta}_{\text{causal}}$). One hundred replicates were used for each scenario.

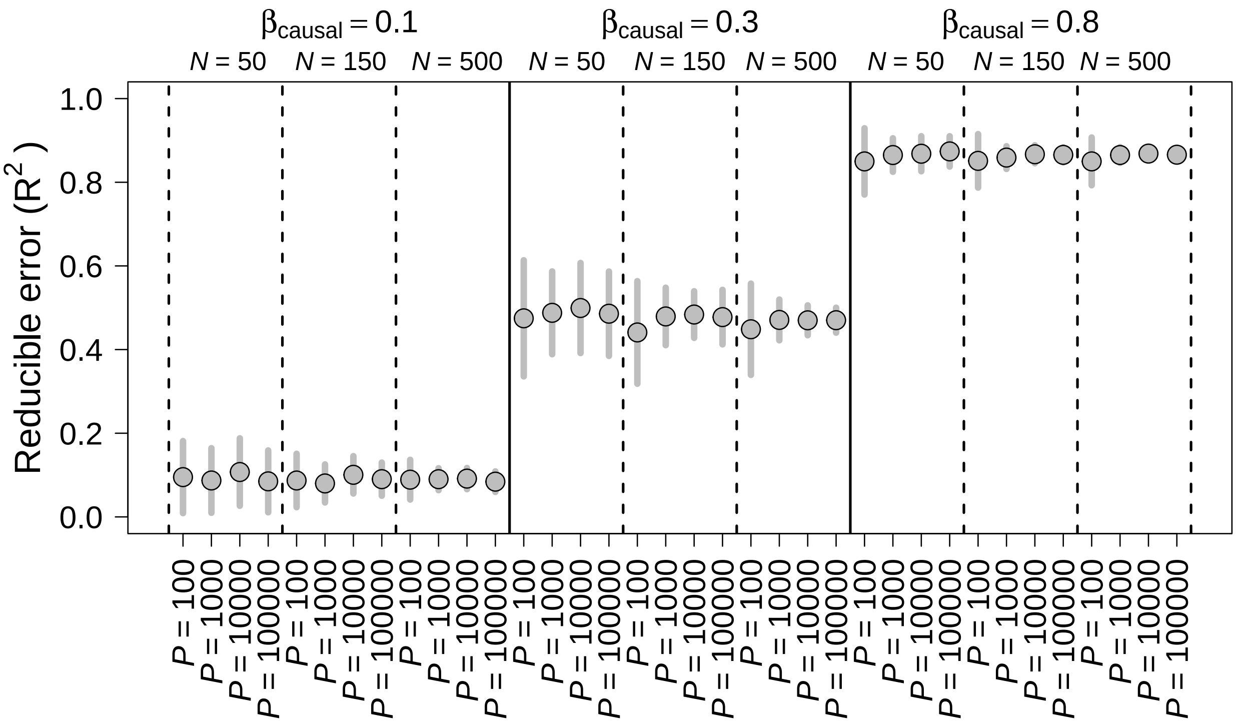

Figure S1: For each core scenario, the reducible error was quantified based on the ability to predict the response using only the 10 causal predictors. Scenarios differed in the number of observations ($N = 50$, 150, or 500), the number of predictors ($P = 100$, 1000, 10000, or 100000), and the effect size of causal predictors ($\boldsymbol{\beta}_{\text{causal}} = 0.1$, 0.3, or 0.8). Median $\mathrm{R}^2$ ($\pm$1sd) from 100 replicates is shown for each scenario.

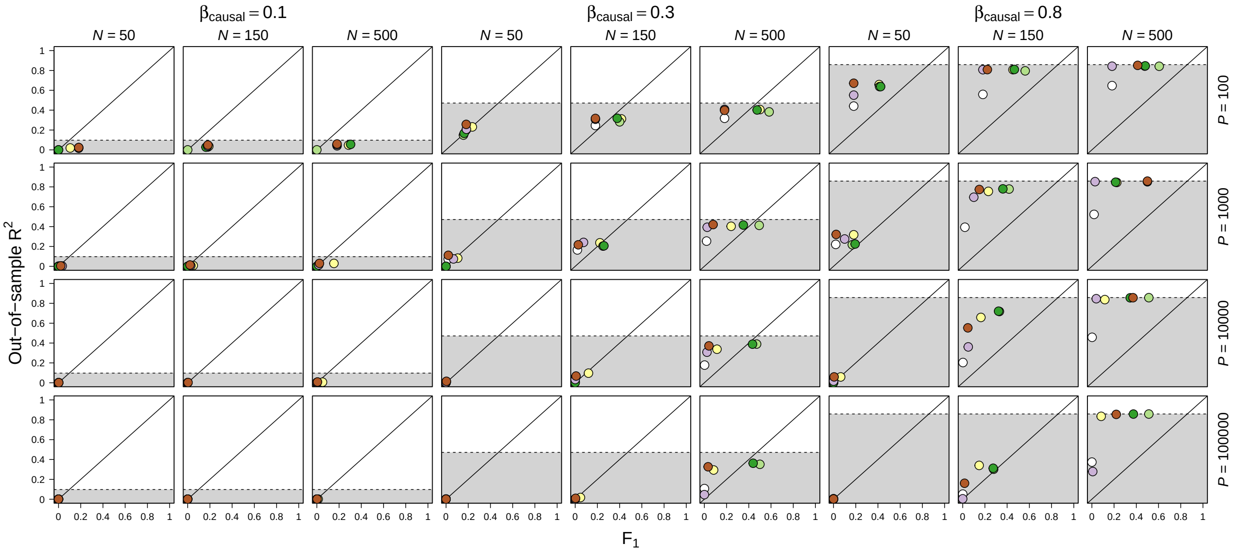

Figure S2: For some scenarios, out-of-sample $R^2$ converged on the target reducible error $R^2$ (shown in gray) even when $F_1$ was relatively low. $F_1$ is the harmonic mean of precision (i.e., the fraction of selected predictors that are truly causal) and sensitivity (i.e., true positive rate): $\frac{2 \times Precision \times Sensitivity}{Precision + Sensitivity}$. $F_1$ was only calculated for analyses that can return truly sparse parameter estimates (i.e., $\boldsymbol{\beta} = 0$; BSLMM, Elastic Net, LASSO, Spike-and-slab) or importance values (Random Forest). Each circle represents the median value from 100 replicate simulations. See Fig. 2 in the main text for color legend.

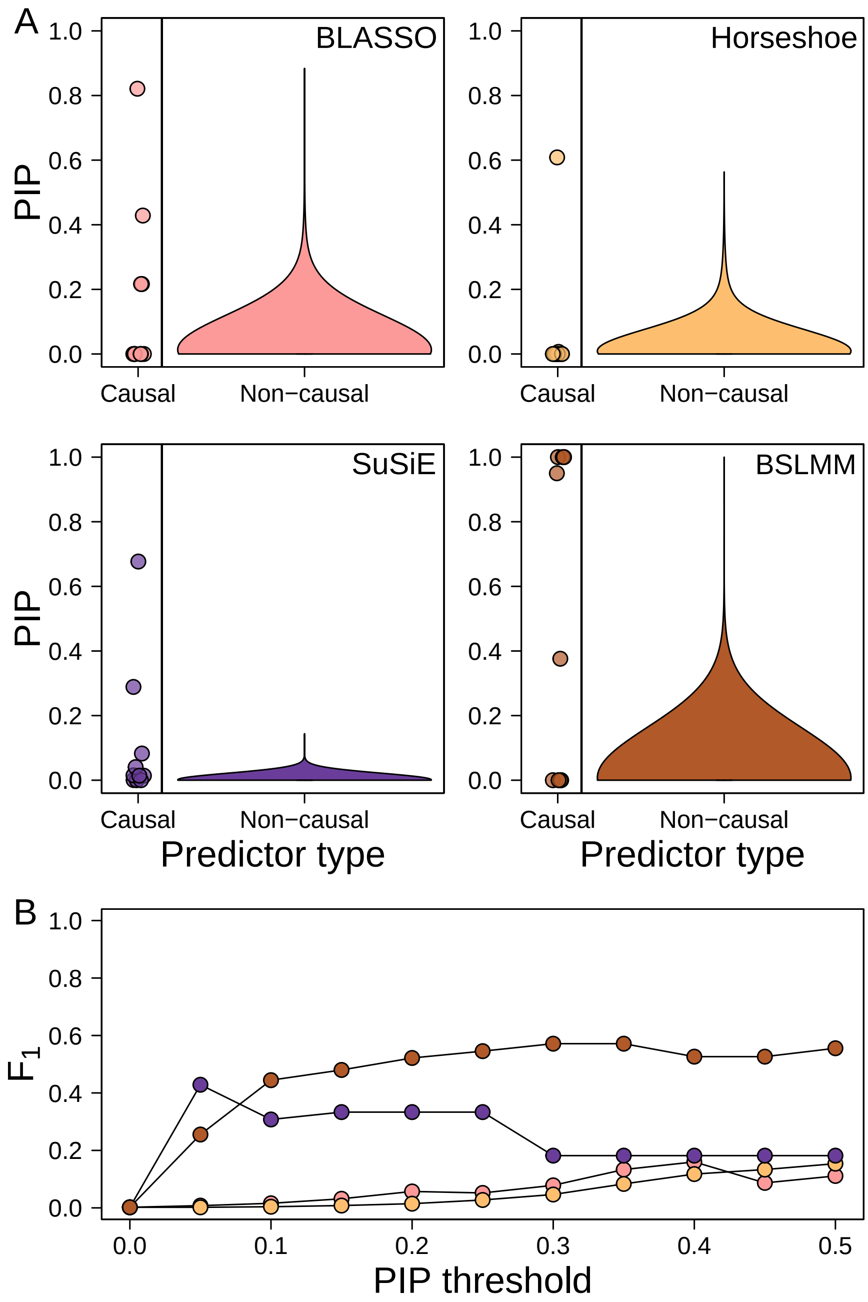

Figure S3: For four Bayesian analyses (BLASSO, Horseshoe, SuSiE, BSLMM), variable selection was also assessed using posterior inclusion probabilities (PIPs) instead of parameter estimates. (A) PIP distributions are shown for the first replicate of scenario 24 (the same example replicate from Fig. 1 in the main text), which had 10 causal predictors ($\boldsymbol{\beta} = 0.8$), 9,990 non-causal predictors ($\boldsymbol{\beta} = 0$), and 150 observations ($N$). (B) Variable selection was evaluated using $F_1$, which is the harmonic mean of precision (i.e., the fraction of selected predictors that are truly causal) and sensitivity (i.e., true positive rate): $\frac{2 \times Precision \times Sensitivity}{Precision + Sensitivity}$. The use of a small PIP threshold of 0.05 (i.e., only predictors with PIP $>$ 0.05 are scored as positives) improved variable selection for BSLMM and SuSiE, whereas larger thresholds were needed to recover more limited gains for BLASSO and Horseshoe.
